# Evidence for plastome loss in the holoparasitic Mystropetalaceae

**DOI:** 10.1101/2025.07.03.662983

**Authors:** Runxian Yu, Matthias Jost, Stefan Wanke, David Bruy, Pan Li, Daniel L. Nickrent, Renchao Zhou

**Affiliations:** School of Life Sciences, State Key Laboratory of Biocontrol and Guangdong Provincial Key Laboratory of Plant Stress Biology, Sun Yat-sen University, Guangzhou 510275, China; State Key Laboratory of Plant Diversity and Specialty Crops, Institute of Botany, The Chinese Academy of Sciences, Beijing 100093, China; University of Chinese Academy of Sciences, Beijing 100049, China; Department of Botany and Molecular Evolution, Senckenberg Research Institute and Natural History Museum, 60325, Frankfurt am Main, Germany; Institute of Ecology, Evolution and Diversity, Goethe-Universitat Frankfurt, 60438, Frankfurt am Main, Germany; Departamento de Botánica, Instituto de Biología, Universidad Nacional Autónoma de México, Mexico City, Mexico; AMAP, Université de Montpellier, IRD, CIRAD, CNRS, INRAE, Montpellier, France; AMAP, IRD, Herbier de Nouvelle-Calédonie, Nouméa, New Caledonia; Laboratory of Systematic & Evolutionary Botany and Biodiversity, College of Life Sciences, Zhejiang University, Hangzhou 310058, China; Plant Biology Section, School of Integrative Plant Science, College of Agriculture and Life Science, Cornell University, Ithaca, NY 14853, USA

**Keywords:** gene loss, mitochondrial genome, parasitic plant, Santalales

## Abstract

Plastome loss is an extremely unusual phenomenon in land plants, even for those that lose their photosynthetic ability. To date, evidence for plastome loss has only been presented for two holoparasitic angiosperm lineages: two genera (*Rafflesia* and *Sapria*) of Rafflesiaceae and one section (*Subulatae*) of *Cuscuta* (Convolvulaceae). Here we propose plastome loss in holoparasitic Mystropetalaceae (Santalales), which consists of three monotypic genera: *Dactylanthus, Hachettea*, and *Mystropetalon*. Using Illumina DNA sequencing data, we successfully assembled the complete mitochondrial genomes of all three genera; however, we failed to recover any plastome sequences from them using reference-based and *de novo* assembly approaches. Illumina RNA sequencing data for *Hachettea* revealed the absence of plastid gene transcripts and losses of many nuclear-encoded genes with plastid function, including genes involved in DNA transcription, translation and protein degradation. Convergent loss of these key genes has also been documented in other plastome-less non-photosynthetic plant lineages. Taken together, we provided compelling evidence for plastome loss in the family Mystropetalaceae.

## Introduction

By definition, holoparasitic plants have lost their ability to photosynthesize, yet nearly all of them have retained a plastome, albeit of reduced size and gene content compared to autotrophic plants (Wicke and Naumann 2018; Sanchez-Puerta et al. 2023). The holoparasitic plant *Pilostyles aethiopica* (Apodanthaceae) has the smallest published plastome of 11.348 kb, containing only five plastid genes unrelated to photosynthesis (Bellot and Renner 2015). To date, only two lineages in land plants have been reported to have lost their plastomes:*Rafflesia* and *Sapria* in Rafflesiaceae (Molina et al. 2014; Cai et al. 2021) and section *Subulatae* of *Cuscuta* in Convolvulaceae (Braukmann et al. 2013; Banerjee and Stefanović 2023), both of which are holoparasitic angiosperms. Another reported case of plastome loss in green plants is from the non-photosynthetic green alga *Polytomella* (Smith and Lee 2014).

The sandalwood order Santalales makes up about half of all parasitic angiosperm species (Nickrent 2020). Two families of this order, Balanophoraceae and Mystropetalaceae, consist of only holoparasitic plants. Known plastomes of Balanophoraceae are highly reduced in size (14.6**–**28.4 kb) and extremely AT-rich in sequence (78.8**–**88.4%), and contain the only two genetic code changes recorded in land plants (Ceriotti et al. 2025). Mystropetalaceae, sister to the mistletoe family Loranthaceae (Su et al. 2015), consists of three monotypic genera, namely, *Dactylanthus, Hachettea*, and *Mystropetalon* (Figure 1), which are endemic to New Zealand, New Caledonia, and South Africa, respectively. To date, no plastid sequences for this family have been deposited in public repositories like GenBank or are available in the literature. There have been a very limited number of mitochondrial and nuclear genes deposited in GenBank for this family. By analyzing Illumina DNA sequencing (DNAseq) data for all three genera of this family and Illumina RNA sequencing (RNAseq) data of *Hachettea* we here provide compelling evidence for plastome loss in the whole family.

**Figure 1.**
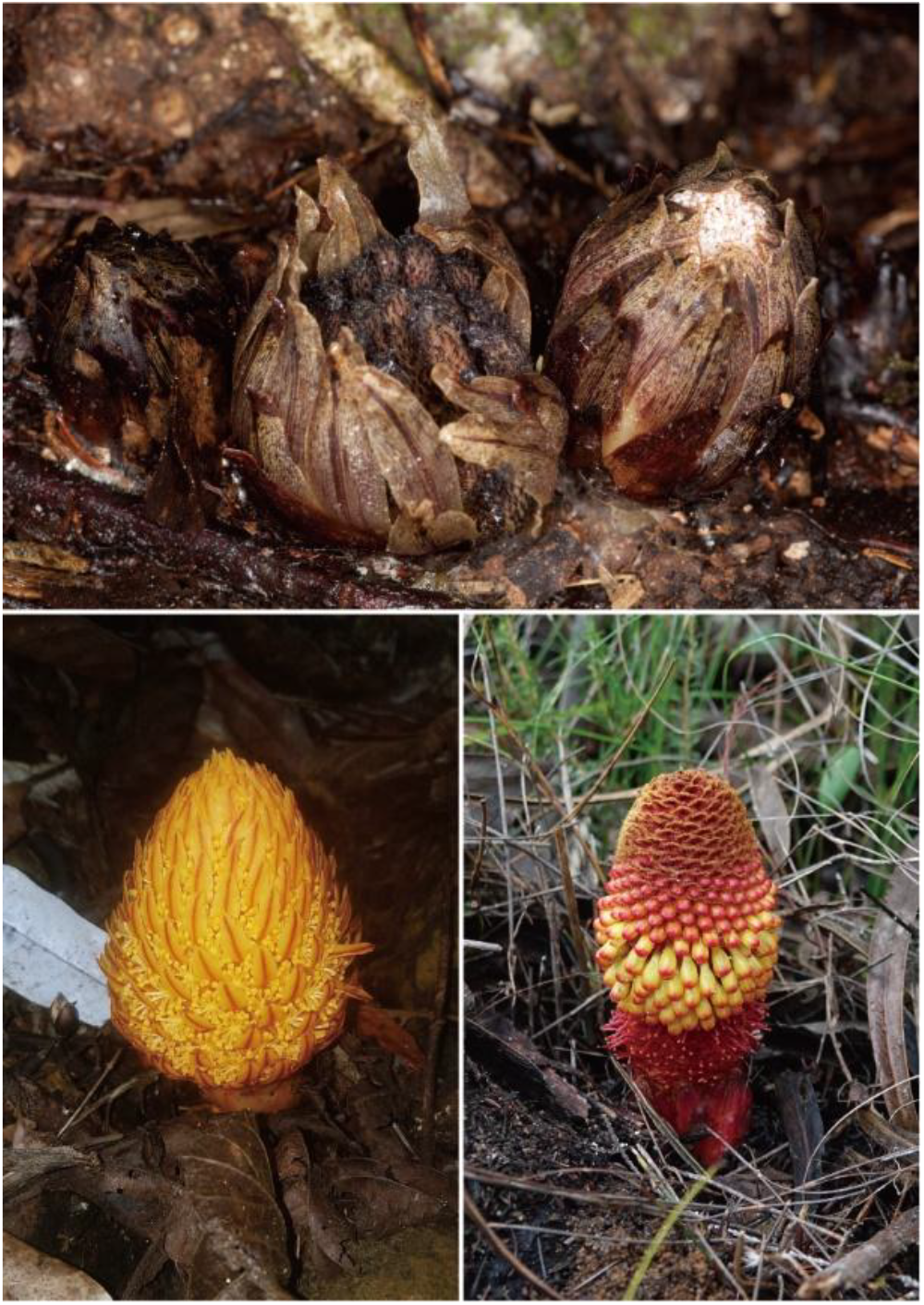
Inflorescences of *Dactylanthus taylori* (top), *Hachettea austrocaledonica* (bottom left) and *Mystropetalon thomii* (bottom right). These photos were provided by Sebastian (Avi) Holzapfel (photo by David Mudge/Nga Manu),David Bruy and Liewen Lin, respectively.

## Results and Discussion

### Mitochondrial genomes (mitogenomes) of Mystropetalaceae contain gene contents typical for most other angiosperms

Using paired-end reads from Illumina DNA sequencing, we successfully assembled the complete mitogenomes of *Dactylanthus taylori, Hachettea austrocaledonica*, and *Mystropetalon thomii*. The mitogenomes of the latter two can be assembled into circular contigs, with their sizes being 750,306 and 526,735 bp, respectively (Figure S1) and average sequencing depths of 98.7 × and 45.2 ×, respectively (Figure S2). The mitogenome of *D. taylori* was assembled into a very complex network consisting of 1,815 contigs (Figure S3), suggesting the presence of numerous repeats. Its total size is 347,672 bp (N50 = 350 bp) with an average sequencing depth of 530.8 ×. The three mitogenomes have GC and gene contents typical for most other flowering plants, except for pseudogenization of *ccmFc* in *H. austrocaledonica* due to an in-frame stop codon reducing the expected coding sequence length by 55 % (Table S1). BLASTN searches of the mitogenomes of *H. austrocaledonica* and *M. thomii* against the plastome of *Erythropalum scandens*, an early-diverging autotroph in Santalales, indicated that some small regions are highly similar to its plastome (Figure 2A; Table S2). Further phylogenetic analysis suggested that several plastid DNA-like regions could derive from ancient intracellular DNA transfers from the plastome or mitochondrion-to-mitochondrion horizontal gene transfers from other plants that had incorporated their own plastid DNA sequences (Figure 2B; Figure S4; Table S2).

**Figure 2.**
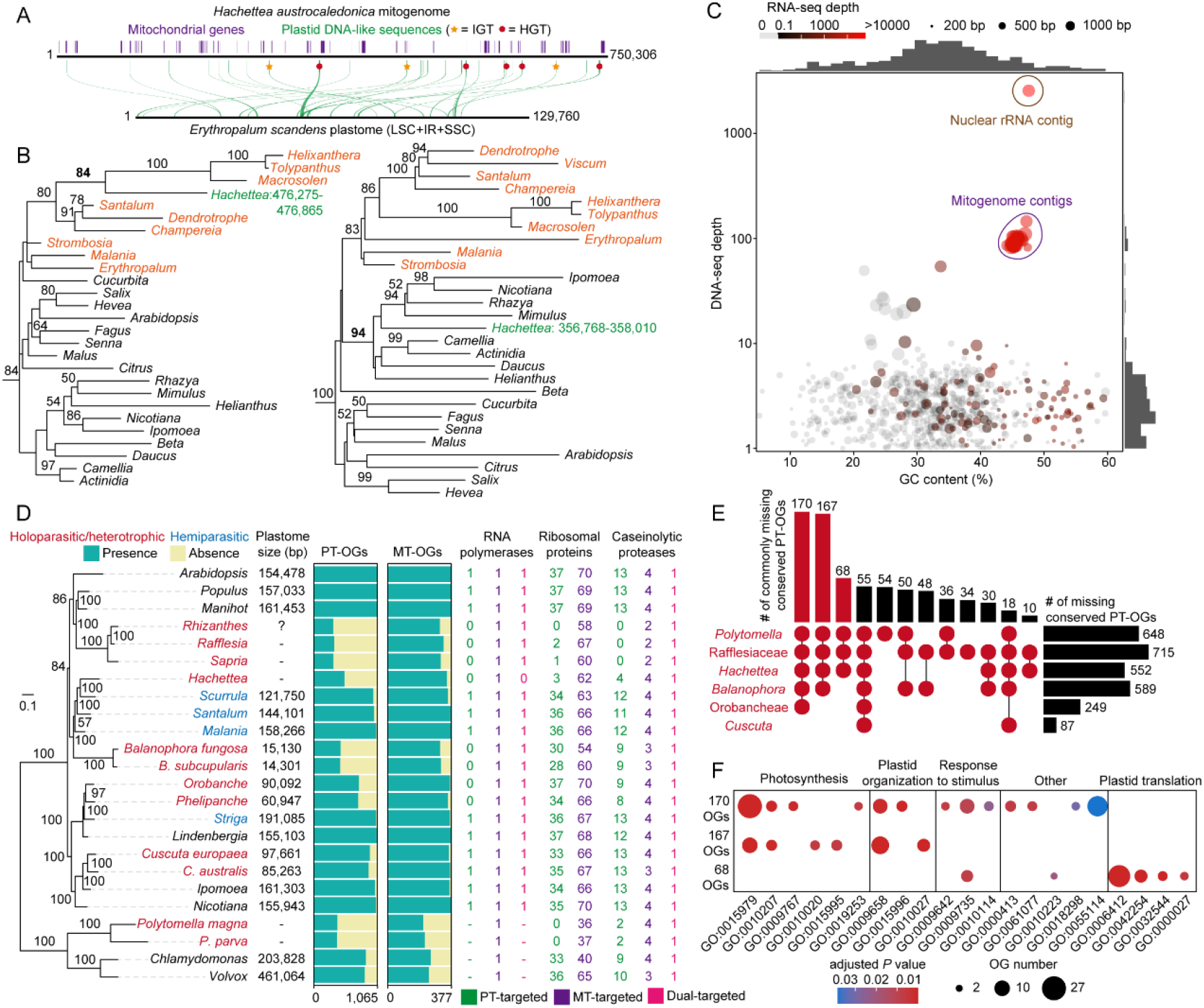
Evidence for plastome loss in the holoparasitic plant *Hachettea austrocaledonica*.(A) Mitochondrial plastid sequences (MtPts) in the mitogenome of *H. austrocaledonica*. MtPts are classified as either from ancient intracellular genic transfer (IGT), horizontal gene transfer (HGT) or unknown origin based on phylogenetic analyses shown in Figure S4. (B) Examples of IGT and HGT-derived MtPts. Sequences of *H. austrocaledonica* (with nucleotide positions in its mitogenome) and other Santalales species are colored in green and orange, respectively. (C) GC contents and sequencing depths for the 879 SPAdes contigs with a BLASTN or NHMMSCAN hit to any of the 12 Santalales plastomes. Mitogenome contigs and a nuclear rRNA gene contig are highlighted. (D) Status of conserved plastid-targeted and mitochondrion-targeted orthogroups in the 24 plants. Phylogenetic relationships of 24 plants are constructed using 221 single-copy genes in the eukaryota_odb10 database in BUSCO. The number of nuclear genes encoding organelle-targeted RNA polymerases, ribosomal proteins and caseinolytic proteases in 24 plants are shown on the right. The symbol “-” means the gene(s) encoding this protein is(are) absent in green algae. PT-OGs: The number of conserved plastid-targeted orthogroups. MT-OGs: The number of conserved mitochondrion-targeted orthogroups. See Table S16 for exact values. (E) An upset plot showing the number of lost conserved plastid-targeted orthogroups among six holoparasitic or non-photosynthetic plants.(F) Gene ontology enrichment analyses for the three largest commonly lost conserved plastid-targeted orthogroups identified in (E).

### No evidence for the presence of plastid contigs in Mystropetalaceae

Plastome assembly using the program GetOrganelle (Jin et al. 2020) with a custom Santalales plastome database (Table S3) as the seed generated 1,227, 49,582, and 67,719 contigs for *D. taylori, H. austrocaledonica*, and *M. thomii*, respectively (Table S4). However, for all three species, contigs meeting the first four criteria for potential plastome contigs proposed in the Methods (< 40 % GC content; having higher DNA sequencing depth than the average sequencing depth of its mitogenome; > 200 bp in length; and having at least one BLASTN or NHMMSCAN hit to a database containing 12 complete plastomes of diverse Santalales) were all identified as either mitochondrial or nuclear rRNAs (Tables S5–S7). Draft genome assembly using the *de novo* assembly program SPAdes (Prjibelski et al. 2020) generated 1,574,389, 12,838,588, and 2,260,724 contigs for *D. taylori, H. austrocaledonica*, and *M. thomii*, respectively (Table S4), among which 123, 879, and 706 contigs have a BLASTN or NHMMSCAN hit to the 12 plastomes of Santalales. However, none of these contigs in any of the three species have a sequencing depth greater than the average sequencing depth of their own mitogenomes, except for mitochondrial or nuclear rRNA sequences (Figure 2C; Figure S5; Tables S8–S10). One 2,780 bp long non-mitochondrial and non-nuclear rRNA gene contig of *D. taylori* (Figure S5; Table S8), with an average sequencing depth of 323.0 × and a GC content of 22.1 %, and containing a 118 bp AT-rich region similar to the plastid *psbC*-*trnN* intergenic spacer of *Tolypanthus maclurei* (Loranthaceae), likely represents a repetitive sequence in the *Dactylanthus* nuclear genome. Additionally, with the exception of mitochondrial and nuclear rRNA gene contigs, all contigs of *H. austrocaledonica* show very low expression levels (Figure 2C), inconsistent with the observation that plant plastid genes typically exhibit very high expression levels (Forsythe et al. 2022). Collectively, no plastome contigs were recovered in the plastome and genome assemblies of the three Mystropetalaceae species.

### No evidence for the presence of plastid gene transcripts in *Hachettea*

Transcriptome assembly of *Hachettea* generated 551,130 transcripts (Table S11), of which 114 have at least one NHMMSCAN hit to any of the 12 Santalales plastomes and both DNAseq depth and RNAseq depth >= 1 ×. Further BLASTN search of the 114 transcripts against sequences of the mitogenome and nuclear rRNA genes of *Hachettea* and the GenBank NT database showed that 64 and six are transcripts of its own mitochondrial genes and nuclear rRNA genes, respectively, and the remaining 44 either are most similar to nuclear rRNA genes (4), nuclear non-rRNA genes (6), mitochondrial DNA (8), plastid DNA of other species (8) or recover no hits (18) (Table S12). Among the eight transcripts most similar to plastid DNA of other species, seven have both very low DNAseq depths (1.0–4.9 ×) and very low RNAseq depths (1.4–5.9 ×), and the remaining one has a slightly higher DNAseq depth (12.5 ×) and a very short hit (88 bp) (Table S12). Such low DNAseq and RNAseq depths of these eight contigs suggests that they are unlikely to be plastid gene transcripts. Therefore, we found no evidence for the presence of plastid gene transcripts in *Hachettea*.

### Substantial loss of nuclear-encoded, plastid-targeted genes in *Hachettea*

Potential plastome loss in *Hachettea* is further supported by substantial loss of nuclear genes that regulate plastid DNA transcription, translation and other functions. We identified orthogroups (OGs) of nuclear genes using the genomes/transcriptome data of 24 plants, including six holoparasitic or non-photosynthetic lineages and their closely related autotrophic and/or hemiparasitic plants (Table S13). The six holo-parasitic or non-photosynthetic lineages show substantial loss of conserved nuclear genes encoding plastid-targeted proteins compared to autotrophic and hemiparasitic plants (Figure 2D; Tables S14– S16). Among the holoparasitic or non-photosynthetic lineages, *Polytomella*, Rafflesiaceae,*Balanophora* and *Hachettea* have lost at least two times more conserved genes encoding plastid-targeted proteins than two other holoparasitic lineages with relatively large plastomes (*Cuscuta* and tribe Orobancheae which includes *Orobanche* and *Phelipanche*) (Figure 2D; Tables S16–S17). Loss of genes encoding mitochondrion-targeted proteins is also observed in the six holoparasitic or non-photosynthetic lineages, although the proportion of loss is smaller than that of plastid-targeted proteins (Figure 2D; Table S16; Table S18). Gene Ontology (GO) enrichment analysis of genes encoding plastid-targeted proteins lost in the six holoparasitic or non-photosynthetic lineages showed that the enriched GO terms for commonly lost plastid-targeted OGs in *Polytomella*, Rafflesiaceae and *Hachettea* were for plastid translation, while those between the three lineages and one or two other holoparasitic lineages were mainly for photosynthesis and plastid organization (Figures 2E and 2F; Table S19).

We further investigated three categories of nuclear-encoded, plastid-targeted proteins related to plastid DNA transcription, plastid translation and plastid protein degradation, and for the purpose of comparison, their counterparts in the mitochondrion (nuclear-encoded, mitochondrion-targeted proteins) (Table S20). The first category comprises nuclear-encoded proteins for plastid DNA transcription. Plastid genes are transcribed by two types of RNA polymerase in angiosperms: plastid-encoded RNA polymerase (PEP) and nuclear-encoded RNA polymerase (NEP), while mitochondrial genes are transcribed by only NEP (Liere et al. 2011). In most land plants, there exists three typical NEPs, namely, plastid-targeted RpoTp, mitochondrion-targeted RpoTm, and dual-targeted RpoTmp (Borner et al. 2015), while in algae there is only mitochondrion-targeted RpoTm, leaving transcription of all plastid genes to PEP (Liere et al. 2011). Among the three genes, both *RpoTp* and *RpoTmp* were found in all eight autotrophic and hemiparasitic flowering plants and one holoparasitic lineage (*Cuscuta*), while only *RpoTmp* was detected in all other holoparasitic flowering plants except *Hachettea* which lacked both *RpoTp* and *RpoTmp* (Figure 2D; Figure S6; Table S20). It is possible that the examined holoparasitic plants with a plastome (except *Cuscuta*) use only *RpoTmp* for plastid DNA transcription. In contrast, the mitochondrion-targeted RpoTm was detected in all 24 examined plants (Figure 2D; Table S20). The absence of both *RpoTp* and *RpoTmp* in *Hachettea* would make plastid DNA transcription highly unlikely.

The second category comprises nuclear-encoded ribosomal proteins for plastid translation. In plastids, translation is performed by a bacterial-type 70S ribosome that is composed of a 50S large subunit and a 30S small subunit. Approximately 75% of all proteins comprising the 50S large subunit (typically 33 different proteins), and 50% of all proteins forming the 30S small subunit (typically 25 different proteins) are encoded by the nuclear genome, while the remaining ribosomal subunits are encoded by the plastome (Sugiura 1995; Tiller and Bock 2014). While almost all 37 nuclear genes encoding these ribosomal subunits were present in the examined 18 plants with a plastome (including holoparasitic plants), none or very few were detected in *Polytomella* (0), *Rafflesia* (2), *Rhizanthes* (0), *Sapria* (1) and *Hachettea* (3) (Figure 2D; Table S20). Mitochondrion-targeted ribosomal protein genes, however, were mostly present in all these lineages (Figure 2D; Table S20). The loss of all or nearly all nuclear genes encoding plastid ribosomal proteins in these lineages makes plastid translation highly unlikely.

The third category involves the plastid caseinolytic protease (CLP) complex, which is vital for maturation of plastid proteins and removal of dysfunctional or unassembled proteins (Nishimura and van Wijk 2015). In *Arabidopsis*, the plastid CLP complex comprises subunits from one plastid-encoded gene, *ClpP1*, and 14 nuclear genes: *ClpP3, ClpP4, ClpP5, ClpP6, ClpR1, ClpR2, ClpR3, ClpR4, ClpC, ClpF, ClpD, ClpT1, ClpT2* and *ClpS1* (Nishimura and van Wijk 2015). With the exception of *Polytomella*, Rafflesiaceae and *Hachettea*, in which 11, 13 and nine of 13 nuclear genes encoding only plastid-targeted proteins were not detected, other species retain at least eight of these genes (Figure 2D; Table S20). All examined species retain *ClpC*, whose product is dually targeted to plastids and mitochondria (Fuchs et al. 2020). Substantial loss of genes encoding the plastid CLP complex can cause dysfunction of translated plastid proteins. In contrast, among the four CLPs targeted only to the mitochondrion (ClpP2, ClpX1, ClpX2 and ClpX3), at least two are retained in all studied species (Figure 2D; Table S20).

### Did Mystropetalaceae species lose their plastid genome?

The lack of any detectable plastid-derived contig in the three genera of Mystropetalaceae, absence of plastid gene transcripts and substantial loss of nuclear genes involved in plastid DNA transcription, translation and protein degradation in *Hachettea*, provides compelling evidence for their plastome loss. Given that the three genera are strongly supported as monophyletic (Su et al. 2015), it is likely that plastome loss happened in the common ancestor of the family. Our study also suggests plastome loss in *Rhizanthes*, due to the same substantial loss of related nuclear genes as *Sapria* and *Rafflesia* (Figure 2D; Table S20). Thus, all three genera of Rafflesiaceae likely have lost their plastomes, implying plastome loss also occurred in their common ancestor.

Despite the frequent occurrence of non-photosynthetic lineages in green plants, plastome-less ones are extremely rare. Why are plastomes retained in nearly all non-photosynthetic plants? The essential tRNA hypothesis states that plastid-encoded *trnE*, besides functioning in translation, is essential for heme biosynthesis in plants, including non-photosynthetic ones (Barbrook et al. 2006). However, the *trnE* gene does not exist in the plastomes of four holoparasitic plant genera: three from Balanophoraceae (*Corynaea, Lophophytum* and *Ombrophytum*; Ceriotti et al. 2025) and one from Apodanthaceae (*Pilostyles*; Bellot and Renner 2015), indicating that the plastid *trnE* gene is not essential for retaining a plastome. We detected 5, 15 and 7 copies of the plastid-*trnE* like gene from genome contigs of the three species of Mystropetalaceae, however, they can clearly be attributed to the nuclear or mitochondrial genomes (Table S21). This could result from ancient IGTs from the plastomes prior to their loss.

Plastids (without their genomes) may be still present in cells of Mystropetalaceae, as was shown in *Polytomella* (Smith and Lee 2014) and *Rafflesia* (Molina et al. 2014). This is supported by the existence of some nuclear-encoded, plastid targeted proteins. For instance, the retention of most genes functioning in heme biosynthesis (nine out of 11) and encoding the TOC-TIC import complex (11 out of 13) in *Hachettea* (Table S22). Ultrastructural studies using transmission electron microscopy may be an avenue to test whether such organelles exist.

## Materials and Methods

### Plant materials and Illumina sequencing

Sampling details for *Dactylanthus taylorii, Hachettea austrocaledonica* and *Mystropetalon thomii* were provided in Table S23. Tissues from one inflorescence of each species were collected. DNA isolation of the three samples and RNA isolation of the *H. austrocaledonica* sample, library construction and Illumina sequencing were conducted following the methods of Yu et al. (2022). Illumina sequencing information is provided in Table S24. The raw Illumina reads were filtered using Trimmomatic v0.39 (Bolger et al. 2014) with default parameters, before further analyses.

### Mitogenome assembly

We used the program GetOrganelle v1.7.1 (Jin et al. 2020) to assemble the mitogenomes of the three species with default parameters. The mitogenomes of *H. austrocaledonica* and *M. thomii* was successfully assembled into circular-mapping contigs, whereas that of *D. taylorii* was assembled into a complex network. To assess the assembly continuity and sequencing depth, for *H. austrocaledonica* and *M. thomii*, we mapped their Illumina DNAseq reads onto their mitogenomes using Bowtie2 v2.5.3 (Langmead et al. 2018) with parameters “--end-to-end --no-mixed --no-discordant”. For *D. taylorii*, we used 52 long gene-containing contigs to calculate its sequencing depth to avoid inaccuracy in sequencing depth assessment due to too many short contigs.

### Plastome assembly

We used two approaches for plastome assembly. First, we used the reference-based program GetOrganelle to assemble their plastomes with the plastomes of 12 Santalales species (Table S3) as the “seed” and -k set to 57, 77, 97. The 12 plastomes included two extremely AT-rich Balanophoraceae plastomes. This was done in case the Mystropetalaceae plastomes have the same highly biased nucleotide composition, which may cause failure in read mapping in the first step of the assembly process. Second, we used the *de novo* assembly program SPAdes v3.13.1 (Prjibelski et al. 2020) to assemble their draft genomes, with the parameter -k set to 57, 77 and 97.

### Nuclear rRNA operon (ETS-18S rRNA-ITS1-5.8S rRNA-ITS2-28S rRNA) assembly

Nuclear rRNA operons of the three species were assembled using GetOrganelle with default parameters except -k set to -k set to 57, 77, 97.

### Identification of potential plastome contigs and plastid *trnE*

Only contigs meeting the following five criteria were considered as potential plastome contigs: 1) < 40 % GC content (Wicke and Naumann 2018), 2) having higher DNA sequencing depth than the average sequencing depth of its mitogenome, 3) > 200 bp in length,4) having at least one BLASTN or NHMMSCAN hit to the 12 plastomes of Santalales and 5) not belonging to mitochondrial sequences or nuclear rRNA operon sequences. The second criterion is based on the observation that plastid DNA generally has much higher sequencing depth than mitochondrial DNA in autotrophic angiosperms (Molina et al. 2014) and three genera of the holoparasitic family Balanophoraceae (Table S25). To identify potential plastome contigs, we first mapped the Illumina DNAseq reads onto the contigs assembled by GetOrganelle and SPAdes, respectively, using Bowtie2 with default parameters except for “--end-to-end”. For *H. austrocaledonica*, we also mapped its RNAseq reads onto its contigs assembled by SPAdes using HISAT2 v2.2.1 (Kim et al. 2019) with default parameters. The mapping results were supplied to SAMtools v1.19.2 (Li et al. 2009) to calculate the average sequencing depth of each contig, with the filtering option -q set to 30. We then identified potential mitochondrial and nuclear rRNA operon contigs by mapping these assembled contigs to their own mitogenomes and nuclear rRNA operons. Contigs with a perfect match were considered as mitochondrial and nuclear rRNA operon contigs. Finally, for the remaining contigs > 200 bp in size, we did a BLASTN search using the 12 plastomes of Santalales as a query (Table S3), with default parameters except -word_size set to 7, and an HMM (Hidden Markov Model)-based search using NHMMSCAN v3.4 (Eddy 2011) with the E-value threshold set to 1e-5. The HMMs were built based on sequence alignments of plastid protein-coding and nuclear rRNA genes of the 12 plastomes of Santalales. To identify potential plastid *trnE* gene sequences in the three species, we did an additional HMM-based search against their own genome contigs > 200 bp using NHMMSCAN. The HMM was built using plastid *trnE* genes of the 12 species of Santalales.

### Mitogenome annotation and identification of mitochondrial plastid sequences

We used the mitogenomes of *Liriodendron tulipifera* (Genbank accession number NC_021152) and seven Santalales species (Table S1) as references to annotate the protein-coding, rRNA and tRNA genes in the Mystropetalaceae mitogenomes. To identify mitochondrial plastid sequences (MtPts) in the mitogenomes of *H. austrocaledonica* and *M. thomii*, we first masked all the annotated mitochondrial genes and then did a BLASTN search against the masked mitogenome using the plastomes of 41 angiosperm species (Table S3) as a query, with default parameters except -word_size set to 7. We did not identify MtPts in the *D. taylorii* mitogenome due to its short contig lengths. To infer the origins of the identified MtPts, phylogenetic analysis was carried out for each MtPt, along with the plastid counterparts of other species. Briefly, nucleotide sequences were aligned using MAFFT v7.505 with the L-INS-i strategy (Nakamura et al. 2018), and a maximum likelihood (ML) tree was inferred in RAxML-NG v1.2.2 (Kozlov et al. 2019), with GTR+Gamma selected as the nucleotide substitution model and 1,000 bootstrap replicates. For a MtPt, if *H. austrocaledonica*/*M. thomii* is nested within Santalales with bootstrap support > 50%, we consider it as a potential intracellular gene transfer (IGT) event; if *H. austrocaledonica*/*M. thomii* is nested within a non-Santalales lineage with bootstrap support > 50%, we consider it as a potential horizontal gene transfer (HGT) event.

### Transcriptome assembly and ORF prediction

A total of 24 species, including 12 holoparasitic or non-photosynthetic plants, four hemiparasitic plants and eight autotrophic plants (Table S13), were used to investigate the loss of nuclear genes encoding organelle-targeted proteins. Although whether the two *Cuscuta* species are hemi- or holoparasites remains debatable, we tentatively regarded them as holoparasites, following the treatments of Cai et al. (2021) and Chen et al. (2023). For species without available genome sequences (Table S11), we *de novo* assembled their transcriptomes using Illumina RNAseq reads with Trinity v2.5.1 (Grabherr et al. 2011) with default parameters. CD-HIT v4.8.1 (Fu et al. 2012) with the parameter “-c 0.9” was used to remove redundancy for the assembled transcripts and then TransDecoder v5.7.1 (https://github.com/TransDecoder/) was employed to predict open reading frames. To eliminate bacterial contamination, contigs were searched against the NCBI bacterial non-redundant database. Contigs with over 80% of aligned length and over 90% identity to bacterial sequences were discarded. The assembly quality was evaluated using the predicted proteins against the eukaryota_odb10 database in BUSCO v5.5.0 (Simão et al. 2015) (Table S26).

### Identification of potential plastid gene transcripts

We used NHMMSCAN to search all assembled transcripts against the HMMs of plastid genes of the 12 Santalales mentioned above, with the E-value threshold set to 1e-5. To determine the sequencing depth for each transcript, we mapped the Illumina DNAseq and RNAseq reads using Bowtie2, setting the parameters “--local” for the former and “--end-to-end --no-mixed --no-discordant” for the latter. Transcripts with at least one NHMMSCAN hit and both DNAseq depth and RNAseq depth >=1 were further searched against sequences of the mitogenome and nuclear rRNA genes of *H. austrocaledonica* and GenBank NT database using BLASTN with default parameters.

### Species tree inference

Of the 255 conserved single-copy genes in the eukaryota_odb10 database in BUSCO, 211 are commonly present in at least seventeen species examined in this study. If a species has two or more copies for a gene, we randomly selected one for phylogenetic analysis. Protein sequences of the 211 genes were aligned using MAFFT with the L-INS-i strategy. Poorly aligned regions were manually trimmed. All aligned sequences were then concatenated into a supermatrix. The best amino acid sequence substitution model (JTT+I+G4) was chosen using ModelTest-NG v0.1.7 (Darriba et al. 2019) and the ML tree was constructed using RAxML-NG with 1,000 bootstrap replicates.

### Ortholog identification

We used Orthofinder v2.5.5 (Emms and Kelly 2019) with default parameters to construct orthogroups (OGs) for all the protein sequences of the 24 species, which generated a total of 56,765 OGs. The largest 100 OGs were removed. When an OG contains proteins of at least eight hemi-parasitic or autotrophic plants, it was defined as a conserved OG. Based on this criterion, 8,795 were considered conserved OGs and used for subsequent analyses (Data S1). We used LOCALIZER v1.0.5 (Sperschneider et al. 2017) with “-p” option, TargetP2 (Armenteros et al. 2019) and DeepLoc v2.0 (Thumuluri et al. 2022) with default parameters to predict the localization of these proteins in the conserved OGs. For each protein, we considered it to be either plastid- or mitochondrion-targeted when there were at least two programs indicating the presence of plastid or mitochondrial signaling peptides with scores or probabilities >=0.5. When one OG contained predicted plastid- or mitochondrion-targeted proteins from at least eight hemi-parasitic or autotrophic plants, it was treated as conserved plastid- or mitochondrion-targeted OG. If a conserved OG doesn’t contain proteins of any examined species from a lineage, this OG is considered as lost in that lineage.

### Analysis of the loss of conserved OGs involved in certain organellar functions

We examined the status of conserved OGs involved in certain organellar functions, including organellar DNA transcription, translation, protein degradation (Table S20). Proteins for these functions and the localizations of these proteins were extracted from the literature (Tiller and Bock 2014; Borner et al. 2015; Nishimura and van Wijk 2015; Woodson et al. 2016; Richardson and Schnell 2019). To infer whether plastids are present or not in the cells of *H. austrocaledonica*, we also examined the OGs related to plastid protein transport (TOC-TIC import complex) and heme biosynthesis (Table S22). *PROTmp* genes of *Scurrula parasitica* var. *graciliflora* and *Ipomoea batatas*, which were not annotated in their gene annotation files, were identified using TBLASTN with default parameters by searching the PROTmp sequence of *Arabidopsis thaliana* (AT5G15700) against their genomes (Table S27). When an OG includes paralogs derived from gene duplication, we constructed a maximum likelihood tree with the same pipeline for species tree construction to distinguish the paralogs (Data S2).

### Gene ontology enrichment analysis

To understand the function of plastid-targeted OGs lost in certain holoparasitic or non-photosynthetic lineages, we carried out a gene ontology (GO) enrichment analysis. Briefly, GO annotation of these OGs was performed based on the biological process (BP)-related GO terms of *A. thaliana* (from the TAIR10 database) or *Chlamydomonas reinhardtii* (from the JGI database). The R package clusterProfiler (Wu et al. 2021) was used to determine significantly enriched GO terms based on a *P* value threshold of 0.05 with a false discovery rate (FDR) correction, using 8,795 conserved OGs as background.

## Acknowledgements

We thank Sebastian (Avi) Holzapfel and Liewen Lin for providing the photos of Mystropetalaceae species. This work was supported financially by the National Natural Science Foundation of China (31811530297) to R.Z.

## Author contributions

RZ, SW, DLN and RY designed research; RZ, RY, MJ, SW, DB, PL and DLN performed research; RY, RZ and MJ analyzed data. RZ, RY, MJ, SW and DLN wrote the manuscript.

## Data availability

All the raw sequencing reads generated in this study were deposited at the National Center for Biotechnology Information (NCBI) under the BioProject PRJNA1268957. All the assembled mitogenomes or mitochondrial genes were submitted to GenBank under accession numbers PV707943, PV742340 and PV843289-PV843329. The assembly graph and annotated coding sequences of newly assembled mitogenomes in this study are also available in the figshare online database (doi: 10.6084/m9.figshare.29246072).

