## Supplemental Figures for "Evidence for plastome loss in the holoparasitic Mystropetalaceae": Supplementary figures 0701.pdf

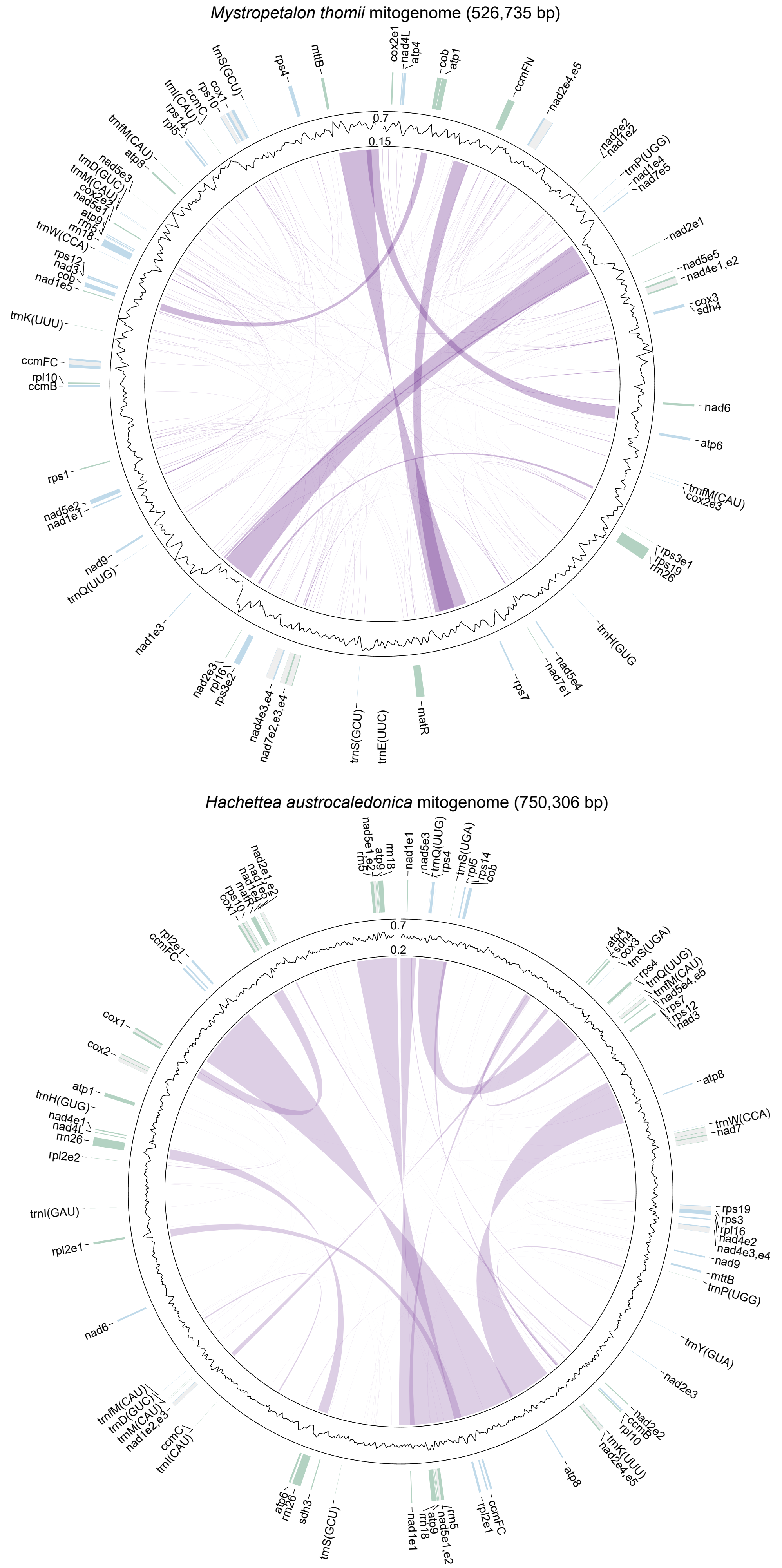


**Figure S1** Mitogenome maps of *Mystropetalon thomii* and *Hachettea austrocaledonica*. Gene annotation is shown in the outer track. Genes colored as blue and green are transcribed clockwise and counter-clockwise, respectively. Grey boxes represent putative *cis*-spliced introns. Line plot displays the GC content across the mitogenomes. Repeats > 50 bp are indicated by purple links.

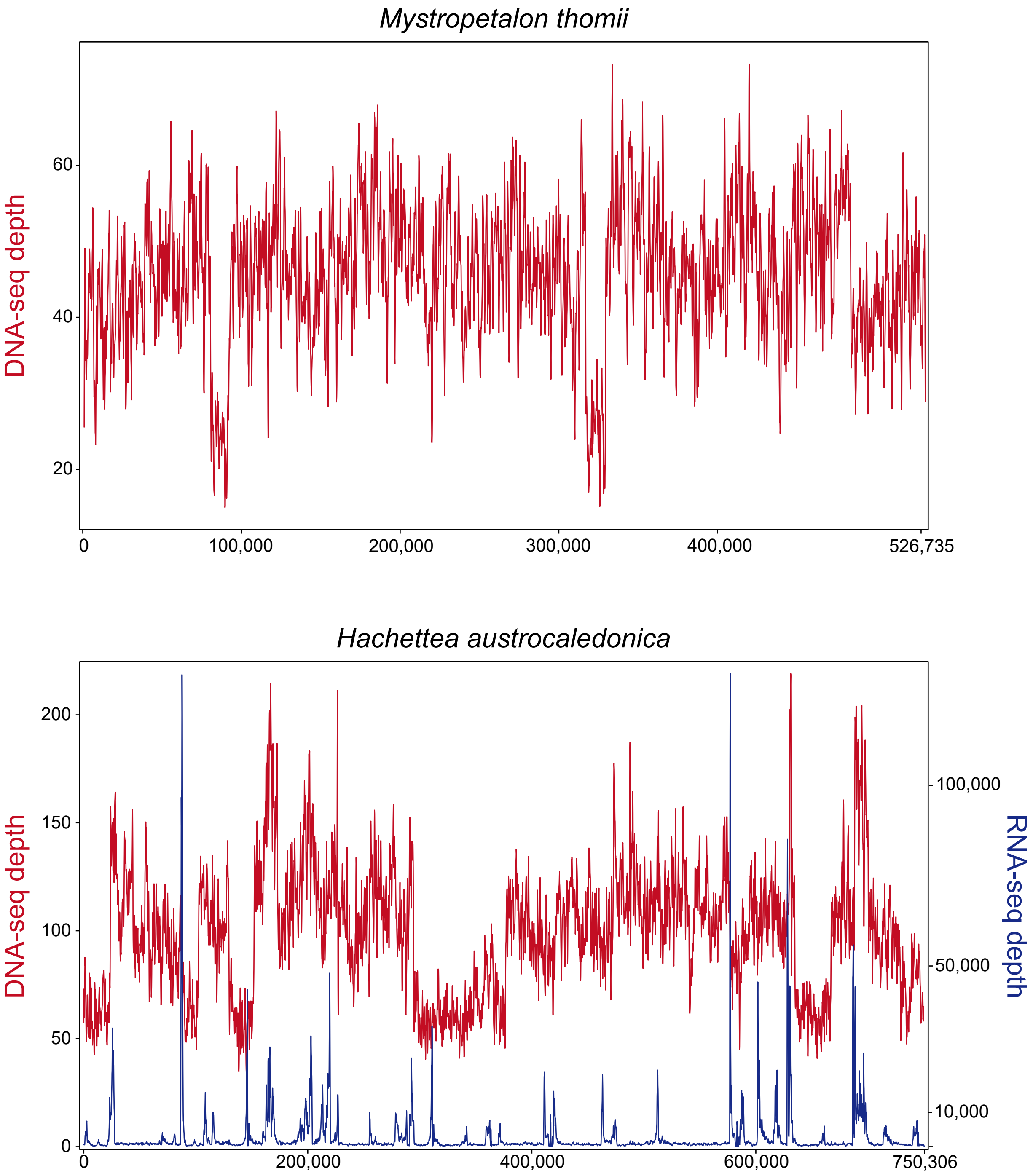


**Figure S2** Sequencing depths of the mitogenomes of *Mystropetalon thomii* and *Hachettea austrocaledonica*.

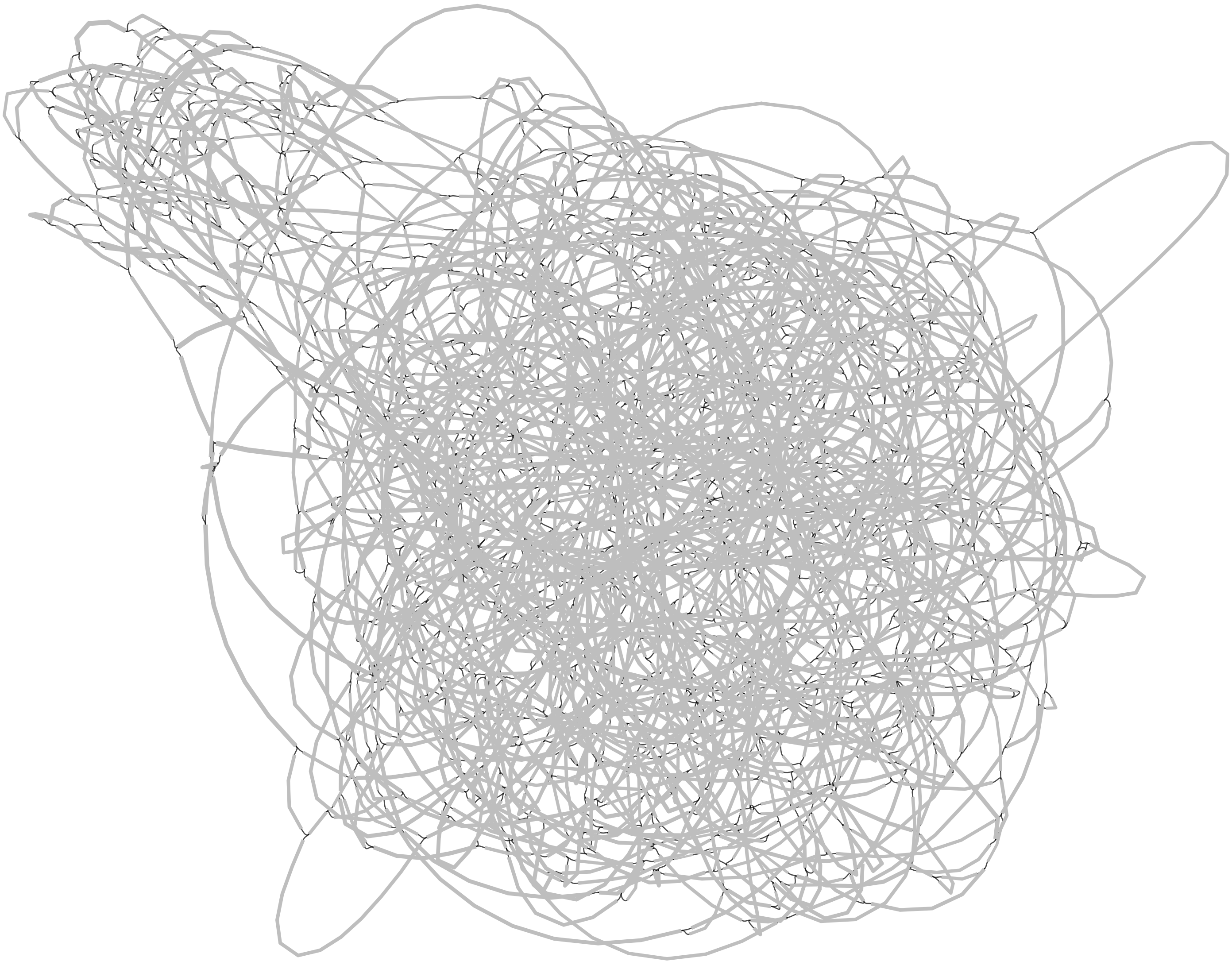

**Figure S3** The assmbly graph of the mitogenome of *Dactylanthus taylori*. Nodes and edges are displayed as gray and black lines. This graph consists of 1,815 contigs with a total size of 347,672 bp (N50 = 350 bp).

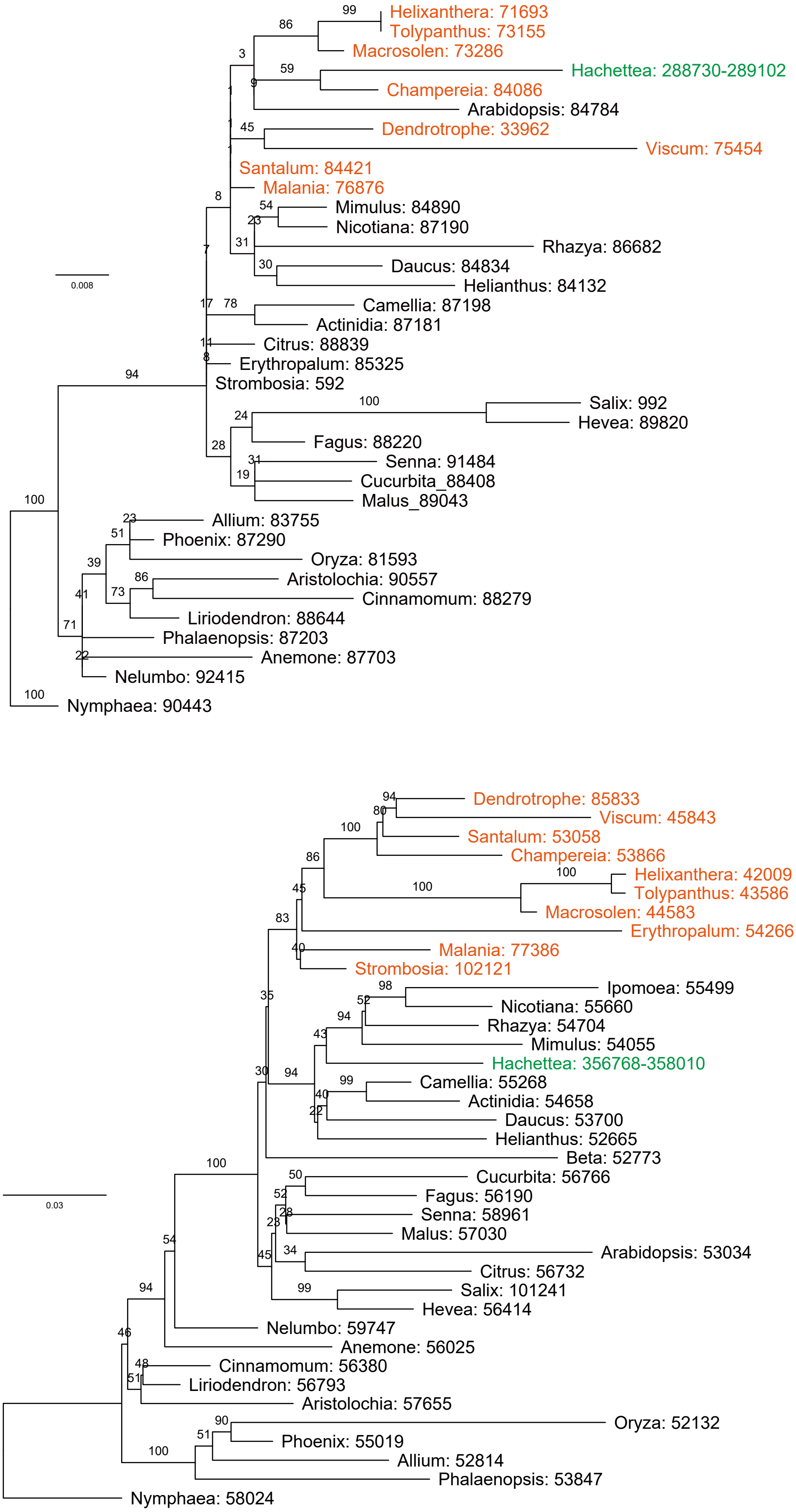


**Figure S4**. Maximum likelihood-based phylogenetic analysis of plastid DNA-like regions in the mitogenomes of *Hachettea austrocaledonica* and *Mystropetalon thomii*. The bootstrap support values are shown on the branch. The nucleotide coordinates (start and end) in the mitogenomes of the two species of Mystropetalaceae and those (only start) in the plastomes of other species are indicated after the genus name. The two species of Mystropetalaceae and other species from Santalales are colored in green and orange, respectively.

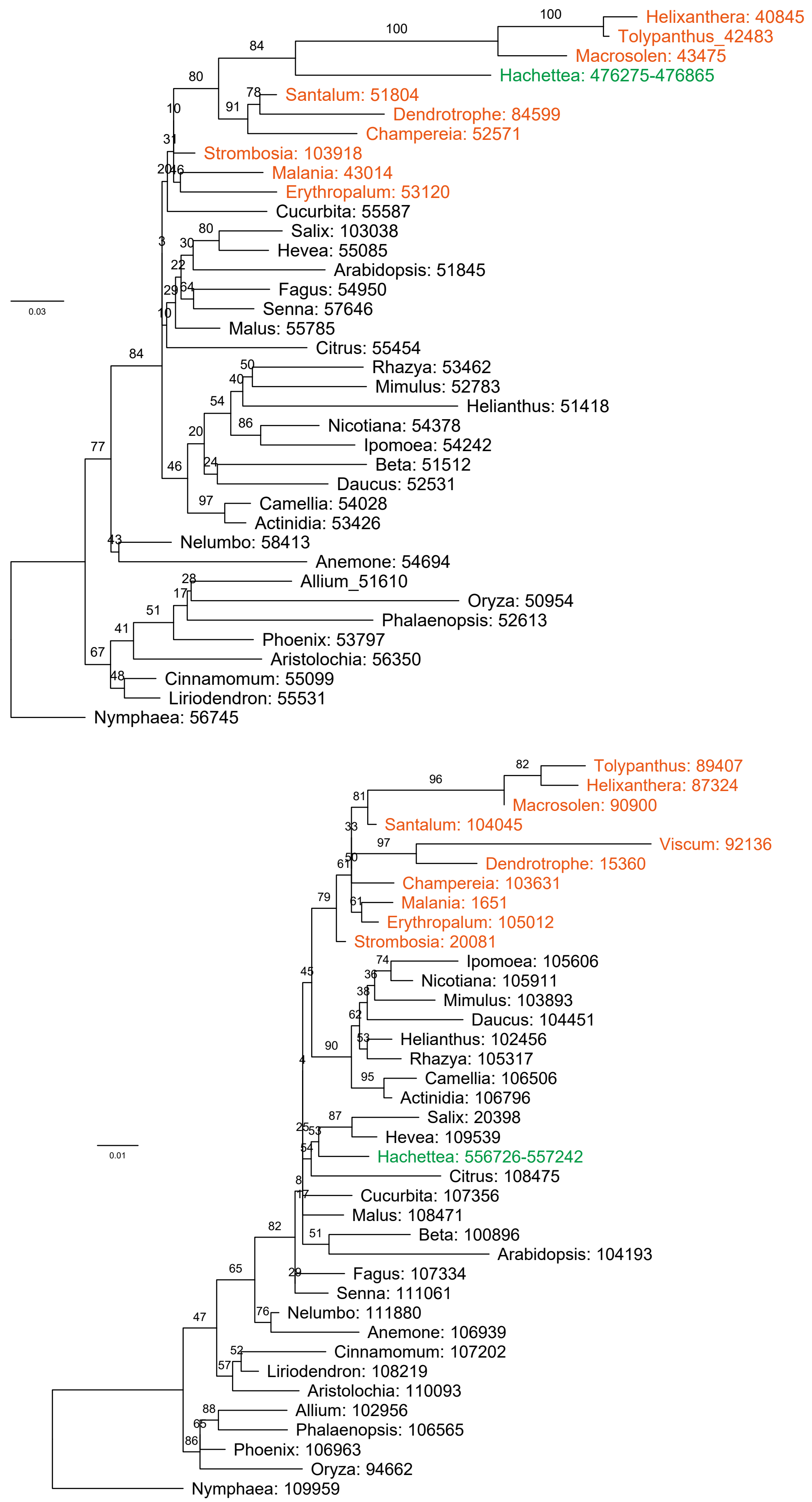


**Figure S4** Maximum likelihood-based phylogenetic analysis of plastid DNA-like regions in the mitogenomes of *Hachettea austrocaledonica* and *Mystropetalon thomii*. The bootstrap support values are shown on the branch. The nucleotide coordinates (start and end) in the mitogenomes of the two species of Mystropetalaceae and those (only start) in the plastomes of other species are indicated after the genus name. The two species of Mystropetalaceae and other species from Santalales are colored in green and orange, respectively.

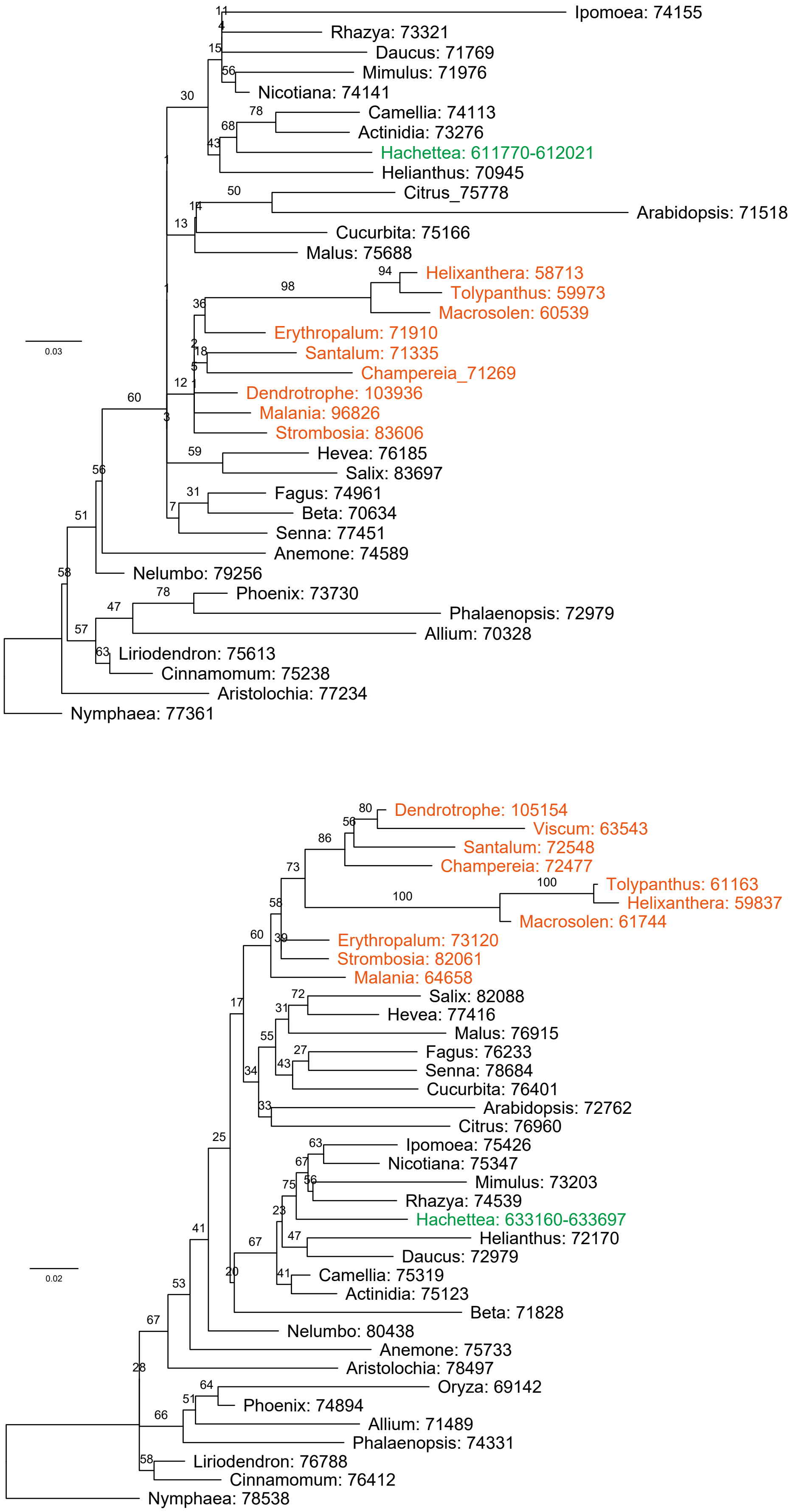


**Figure S4** Maximum likelihood-based phylogenetic analysis of plastid DNA-like regions in the mitogenomes of *Hachettea austrocaledonica* and *Mystropetalon thomii*. The bootstrap support values are shown on the branch. The nucleotide coordinates (start and end) in the mitogenomes of the two species of Mystropetalaceae and those (only start) in the plastomes of other species are indicated after the genus name. The two species of Mystropetalaceae and other species from Santalales are colored in green and orange, respectively.

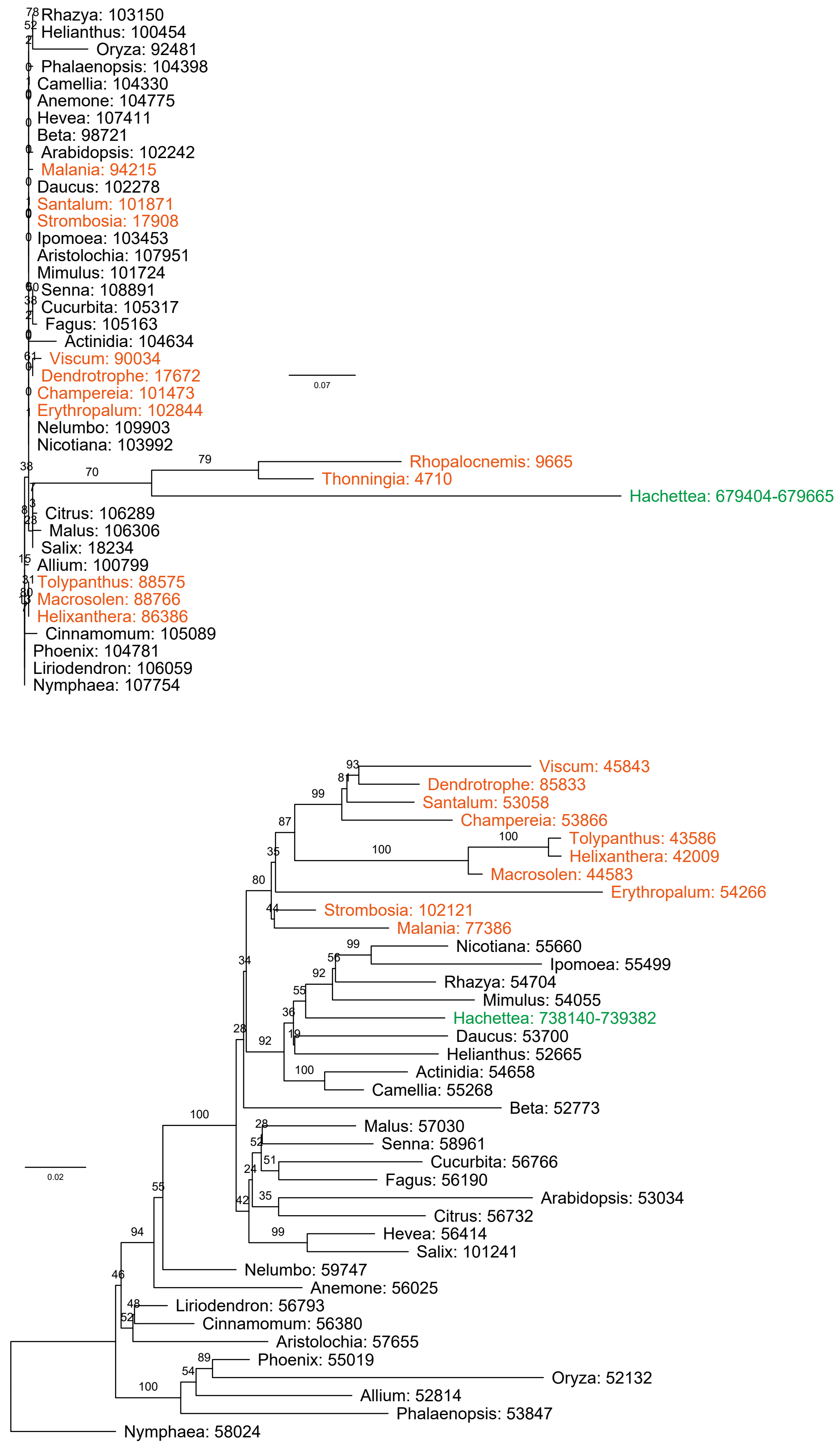


**Figure S4** Maximum likelihood-based phylogenetic analysis of plastid DNA-like regions in the mitogenomes of *Hachettea austrocaledonica* and *Mystropetalon thomii*. The bootstrap support values are shown on the branch. The nucleotide coordinates (start and end) in the mitogenomes of the two species of Mystropetalaceae and those (only start) in the plastomes of other species are indicated after the genus name. The two species of Mystropetalaceae and other species from Santalales are colored in green and orange, respectively.

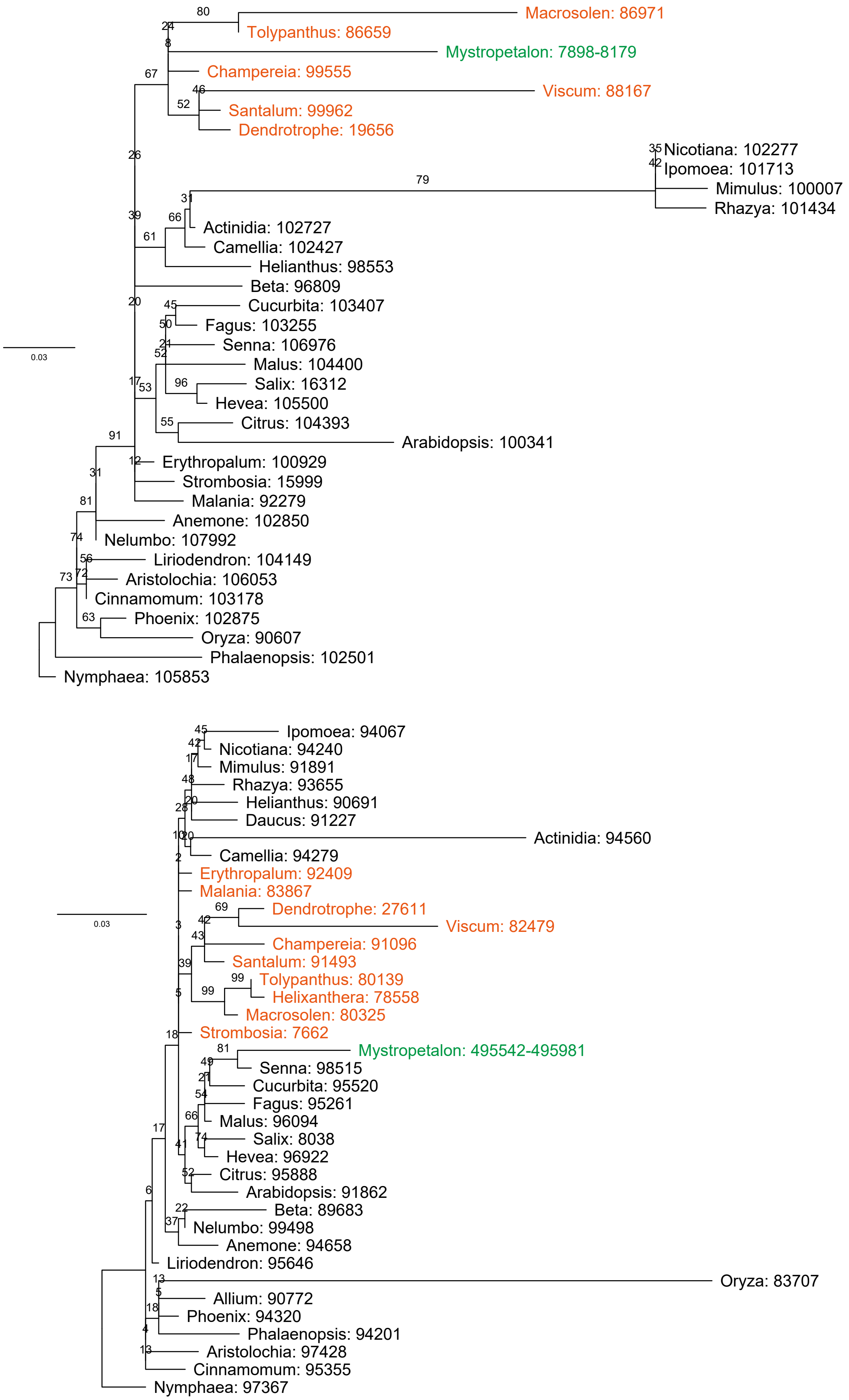


**Figure S4** Maximum likelihood-based phylogenetic analysis of plastid DNA-like regions in the mitogenomes of *Hachettea austrocaledonica* and *Mystropetalon thomii*. The bootstrap support values are shown on the branch. The nucleotide coordinates (start and end) in the mitogenomes of the two species of Mystropetalaceae and those (only start) in the plastomes of other species are indicated after the genus name. The two species of Mystropetalaceae and other species from Santalales are colored in green and orange, respectively.

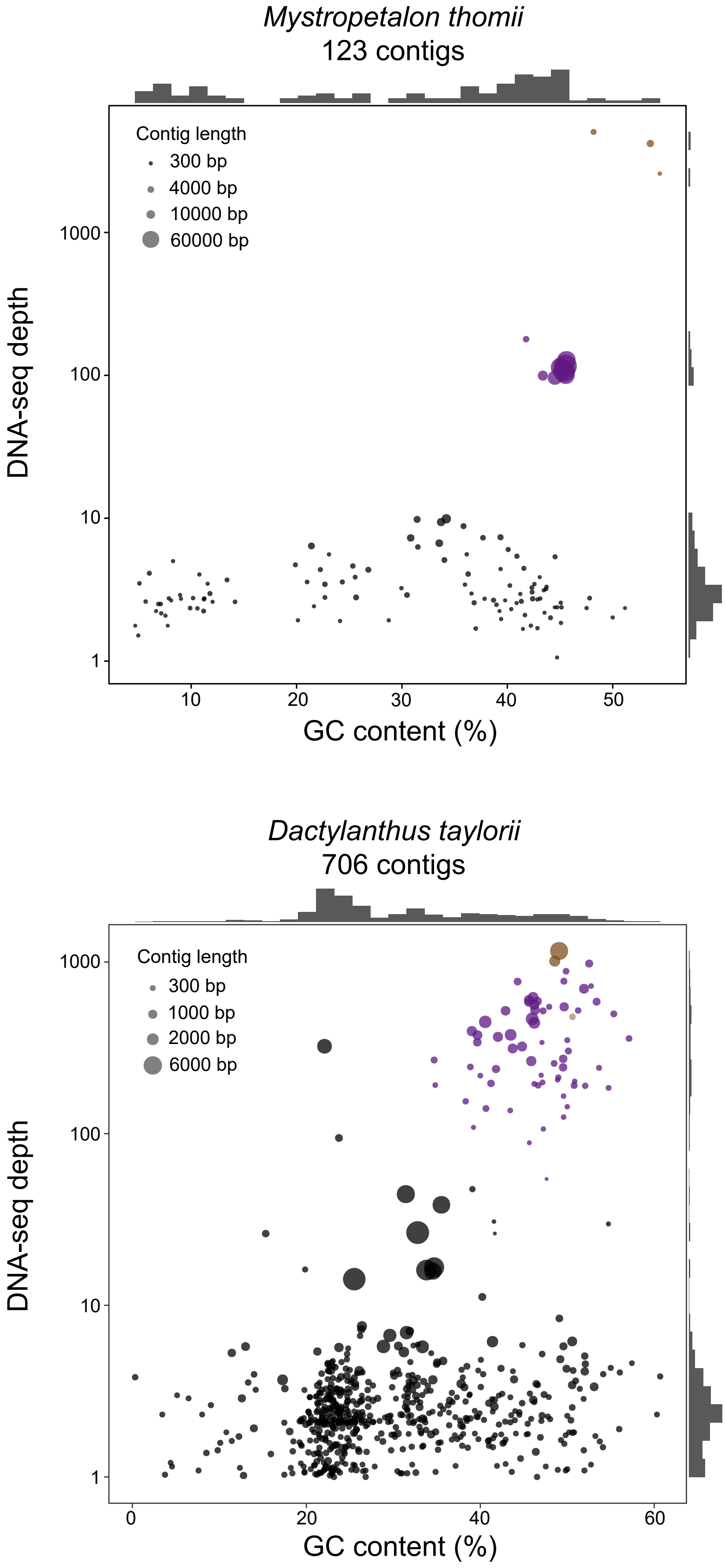


**Figure S5** GC contents and DNA sequencing depths for SPAdes contigs of *Mystropetalon thomii* and *Dactylanthus taylorii* with a BLASTN or/and HMMSCAN hit to the plastomes of 12 Santalales species. Mitogenome and nuclear rRNA contigs are colored in purple and brown, respectively.

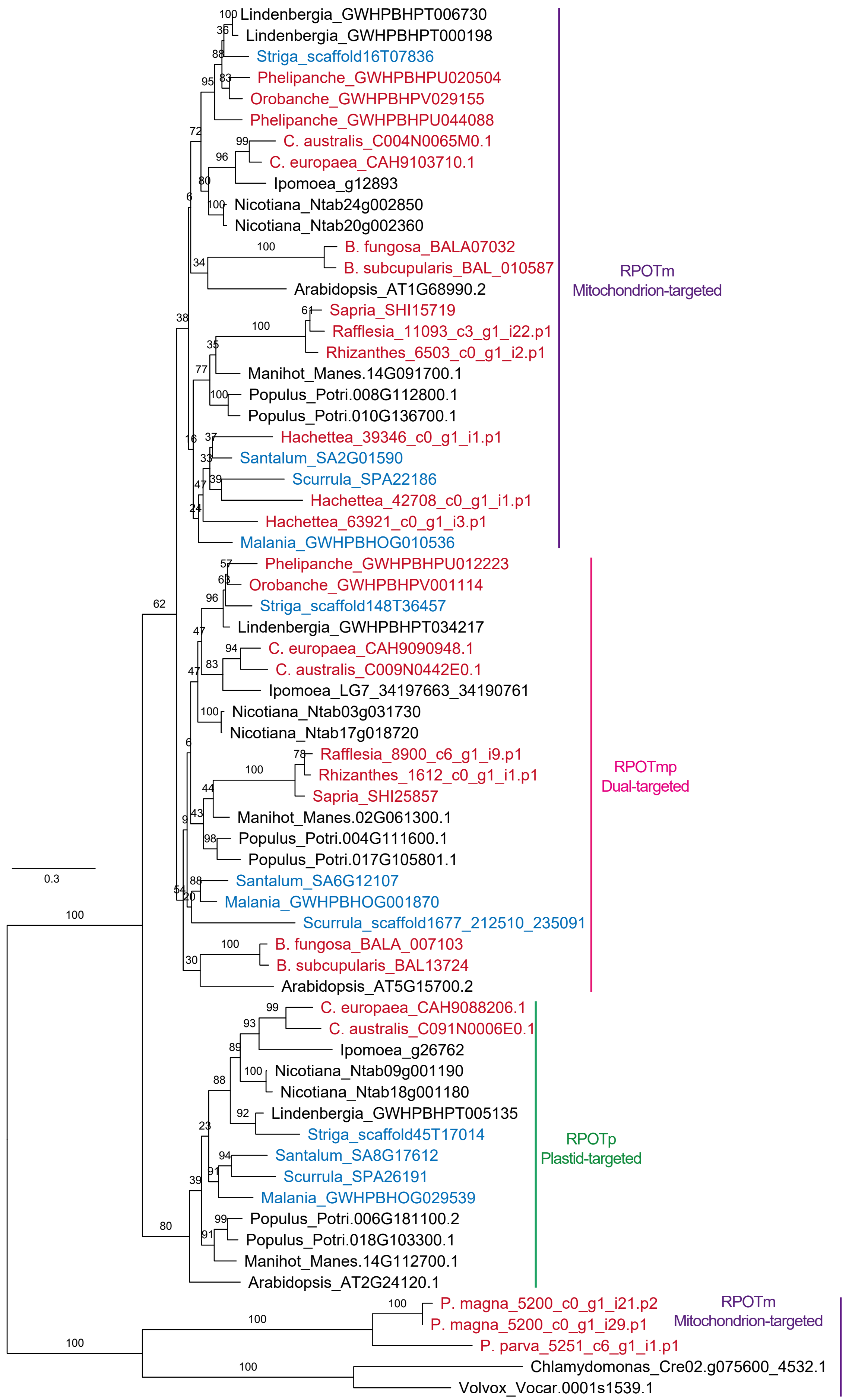


**Figure S6** A maximum likelihood tree of 24 plants based on protein sequences of nuclear-encoded organelle-targeted RNA polymerases. Holoparasitic/non-photosynthetic and hemiparasitic plants are colored in red and blue, respectively.
